# Adaptive laboratory evolution unlocks membrane permeability as a key limitation in long-chain alcohol metabolism by *Pseudomonas putida* KT2440

**DOI:** 10.64898/2026.01.19.700371

**Authors:** Raúl Mireles, Lianet Noda-García

## Abstract

*Pseudomonas putida* KT2440, renowned for its diverse metabolic capabilities, is a promising platform for downstream processing and revalorization of recalcitrant molecules. In this study, we examined and optimized *P. putida* KT2440’s ability to utilize long-chain alcohols. These molecules are byproducts of the degradation of polyethylene (PE), the most widely used plastic. Using them as feedstock for microbial growth would close the plastic-derived carbon cycle, reducing environmental pollution. First, we discovered that *P. putida* KT2440 can use long-chain alcohols as the sole carbon and energy source. Using adaptive laboratory evolution (ALE), we generated variants with improved growth rates on long-chain alcohols, specifically 1-hexadecanol and 1-eicosanol. Mutations that became fixed during ALE provided insights into the mechanism, highlighting the importance of cell-substrate interaction. By heterologously expressing a hydrocarbon transporter-encoding gene, we successfully reproduced the ALE-derived phenotype, demonstrating that the bottleneck in long-chain alcohol utilization is not substrate transformation but uptake. These findings lay the groundwork for the potential application of *P. putida* KT2440 for the degradation of PE.

## 1. Introduction

Plastics are an integral part of contemporary life (Geyer, 2017). Their low production costs and versatile functionality made them ubiquitous across all economic sectors (e.g., from the food industry to healthcare) (OECD, 2022a). This demand has driven global plastics production to an average of ∼400 million metric tons (Mt) per year over the last five years, a market that is projected to produce 1,231 Mt by 2060 (PlasticsEurope, 2025, OECD, 2022a). The increasing demand is accompanied by inadequate disposal. Post-use of this waste is mostly landfilled (46%), dumped (22%), or incinerated (17%), with only 15% collected for recycling (OECD, 2022a). These shares highlight a persistent gap between consumption and end-of-life capacity to fulfil the circularity of plastics.

Among plastics, polyethylene (PE) is the main contributor to production (110 Mt, 24%) and waste (93.9 Mt, 26.6%) (OECD, 2022b). PE is an oil-derived polymer with strong C-C and C-H bonds, making it highly resistant to degradation (Schwab, 2024). Upon environmental discharge, PE films have an estimated half-life of 2,500 years (Chamas, 2020). While the impact of discarded PE in the environment is yet to be fully understood, evidence points to greenhouse-gas emissions from weathering, release of dissolved organic carbon that shifts carbon cycling, vectoring of pollutants, and physical smothering that restricts light and oxygen, degrading habitat quality (Royer, 2018; Romera-Castillo, 2018; Koelmans, 2016; Greens; 2015). Thus, proper disposal of PE is of paramount importance.

PE recycling efforts have focused on developing sustainable strategies to repurpose PE by breaking it down into simpler molecules (Bui, 2023). The conversion of PE yields more manageable oxygenated low-molecular-weight compounds, including long-chain aliphatic alcohols (i.e., fatty alcohols) such as 1-hexadecanol (C16OH) and 1-eicosanol (C20OH) (Hakkarainen and Albertsson, 2004; Bui, 2023i). Given the abundance of PE production and low-cost, fatty alcohols derived from PE treatment are an attractive feedstock for microbial growth. Their utilization in fermentation processes will thereby close the PE-carbon cycle sustainably.

*Pseudomonas putida* KT2440, a soil bacterium renowned for its metabolic versatility, robustness, and genetic tractability, has emerged as a leading platform for biotechnological applications (Weimer, 2020; Martínez-García, 2014, Nickel, 2018). Previous research has demonstrated the organism’s ability to metabolize a wide range of aliphatic alcohols, including short- (e.g., ethanol, C2OH) and medium-chain (e.g., 1-dodecanol, C12OH), indicating its catabolic potential for fatty alcohol substrates relevant to PE degradation pathways (Thompson, 2020; Lu, 2023).

Here, we examined *P. putida* KT2440’s ability to grow on 1-hexadecanol (C16OH) and 1-eicosanol (C20OH) as carbon sources. We found that it can utilize these molecules as a sole carbon and energy source, thereby expanding the known repertoire of recalcitrant molecules that this versatile bacterium already metabolizes. Furthermore, we performed five cycles of adaptive laboratory evolution (ALE) to generate variants with enhanced capacity to metabolize these molecules. In this short time, the isolated adapted variants grew 1.4 (±0.1) times faster than the wild-type strain. Comparative genomic analysis of wild-type versus adapted variants, along with experimental validation, revealed that substrate uptake, rather than metabolic processing itself, constitutes the primary bottleneck for the efficient utilization of fatty alcohols. This work enhances the fundamental understanding of fatty alcohol metabolism in *P. putida* KT2440 and provides a foundation for engineering microbial platforms capable of valorizing plastic waste into valuable products, contributing to circular economy initiatives.

## 2. Results

### 2.1 Pseudomonas putida KT2440 can utilize fatty alcohols as a sole carbon source

Until now, the utilization of aliphatic alcohols by wild-type *Pseudomonas putida* KT2440 has been described only for short- and medium-chain alcohols up to 12 carbon atoms in the main chain (e.g., ethanol, C2OH to 1-dodecanol, C12OH) (Thompson, 2020; Lu, 2023). Given the limited knowledge of its capacity to metabolize longer molecules, we tested whether wild-type *P. putida* KT2440 could utilize 1-hexadecanol (C16OH) and 1-eicosanol (C20OH) as the sole carbon source. We tested three concentrations (2.5, 5.0, and 10.0 mM) to assess concentration-dependent effects on growth (Figure 1A–B). Growth rates and carrying capacity exhibited a clear dependence on substrate concentration, with higher concentrations supporting the fastest growth rates and biomass accumulation for both fatty alcohols (Figure 1C–F). These findings indicate that at these concentrations, the low solubility of fatty alcohols does not appear to be a bottleneck for utilization and that the substrates are not toxic, as observed with other alcohol substrates like 1-octanol (Huesemann, 2004; Kongpol, 2014).

**Figure 1.**
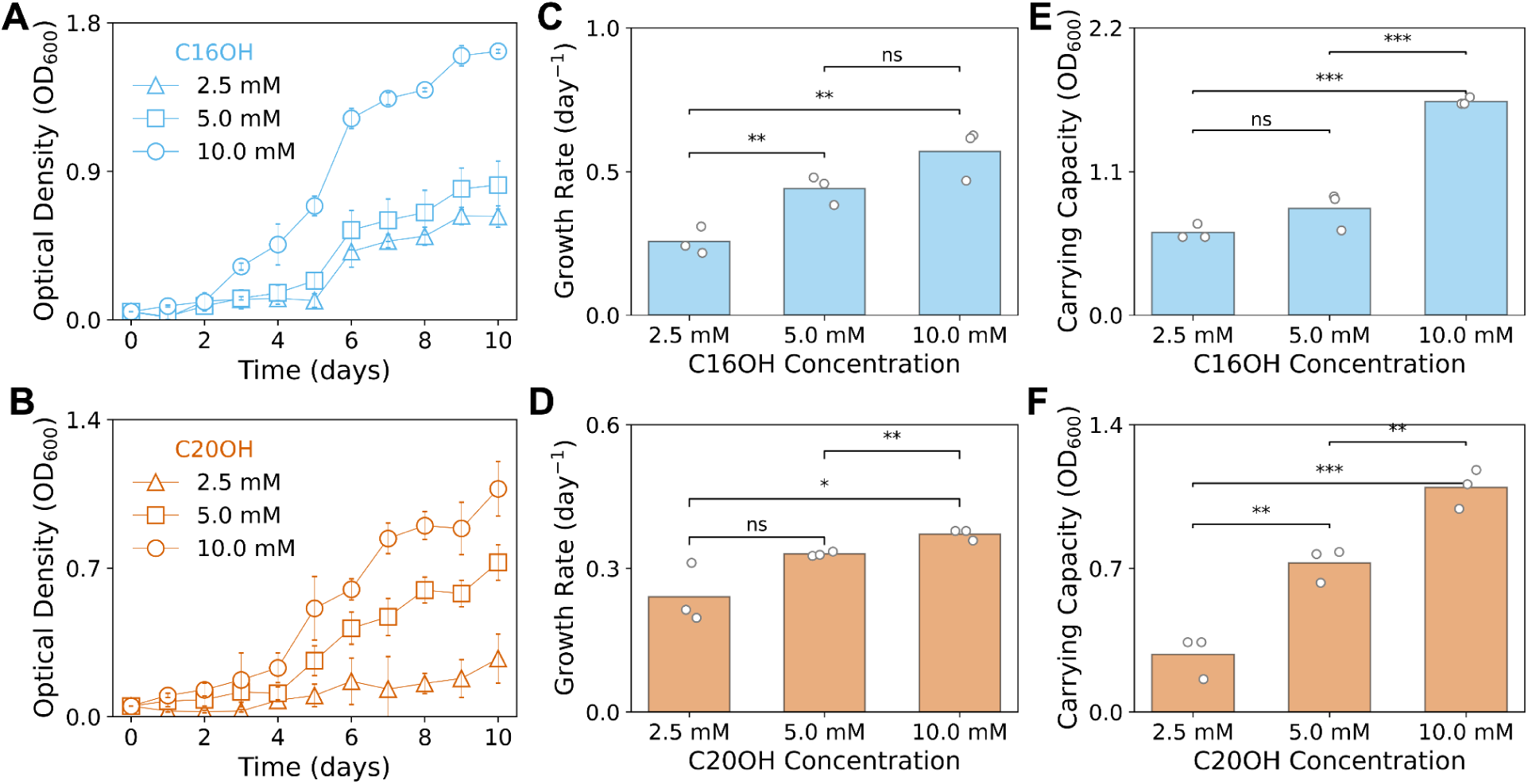
*Pseudomonas putida* KT2440 uses long-chain alcohols as the sole carbon source. Growth curves of wild-type *P. putida* KT2440 on varying concentrations of C16OH (A) and C20OH (B). Growth rate (C, D) and carrying capacity (E, F) for the tested concentrations. Error bars represent the standard deviation of three biological replicates. Individual data points are shown. Statistical significance was assessed using t-tests; *p* < 0.05 (*), *p* < 0.01 (**), and not significant (ns).

### 2.2 Adaptive laboratory evolution enhances wild-type Pseudomonas putida KT2440 growth rate on fatty alcohols

Having established that *P. putida* KT2440 can use fatty alcohols as a carbon and energy source, we sought to identify the bottleneck in their utilization to guide future engineering efforts toward optimization. We performed five passages of adaptive laboratory evolution (ALE) using 1-hexadecanol (C16OH) and 1-eicosanol (C20OH) as the sole carbon and energy source at 2.5 mM (Figure 2A). We initiated three independent evolution lines for each substrate starting from wild-type *P. putida* KT2440. Initially, we grew the wild-type strain for 23 days in C16OH and for 25 days in C20OH. Cultures were then diluted into fresh minimal media (initial optical density at 600 nm, OD_600_, 0.05) containing the same carbon and energy source and maintaining the same concentration. For the next four passages, this dilution was performed when the OD_600_ remained unchanged over two consecutive days, indicating that the culture had entered the stationary phase. We observed homogeneous, rapid phenotypic optimization immediately after the first passage, which remained consistent across the subsequent four passages. After passage one, dilution times decreased from 23 and 25 days to 7 and 10 days and remained constant throughout the rest of the ALE process (Figure S1).

**Figure 2.**
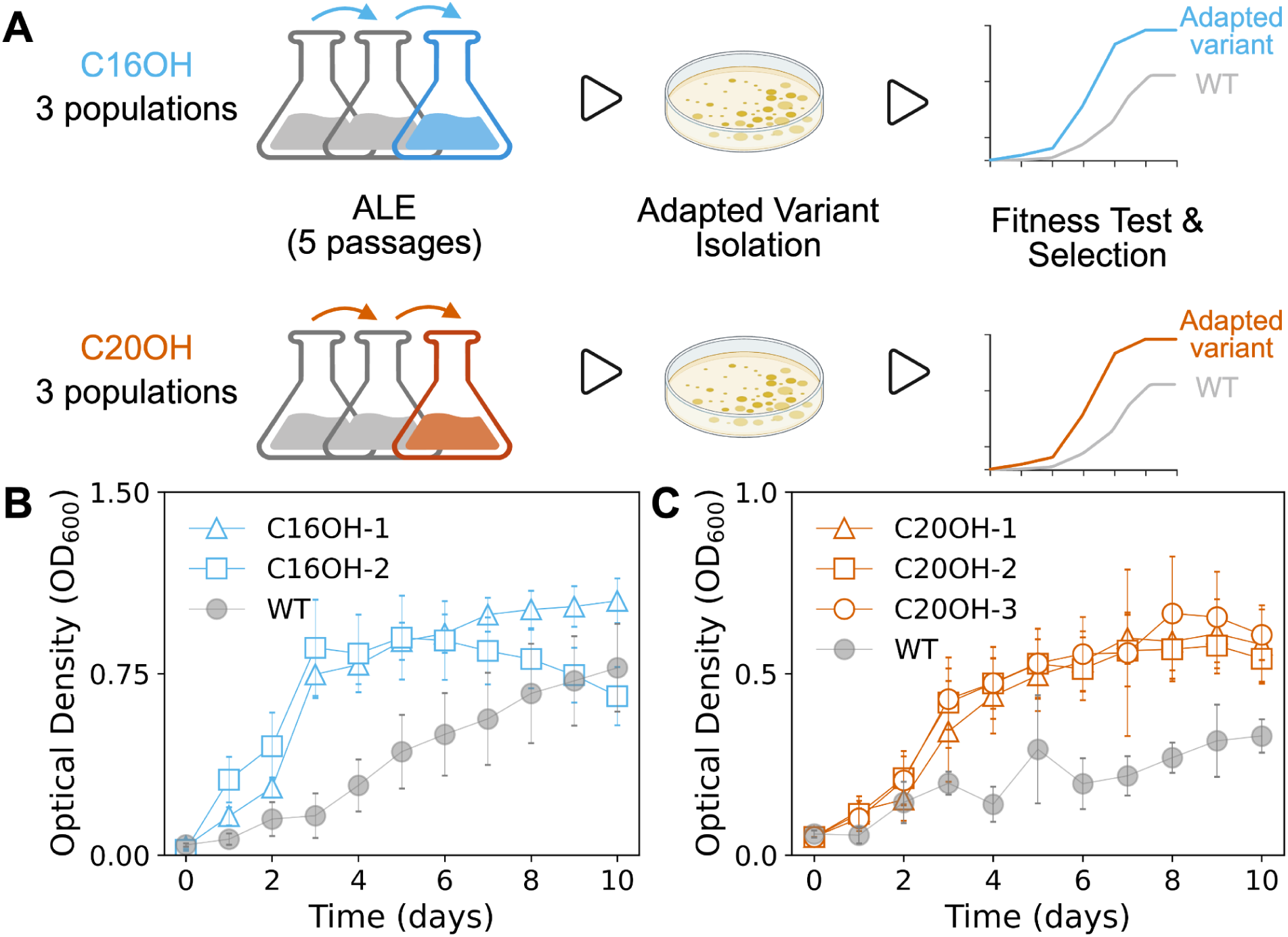
Adaptive laboratory evolution (ALE) of *Pseudomonas putida* KT2440 on long-chain alcohol enhances the growth rate of adapted variants. (A) Schematic overview of the ALE workflow. Growth curves of selected adapted variants from the ALE experiment vs. wild-type (WT) on 2.5 mM C16OH (B) and 2.5 mM C20OH (C). Error bars indicate standard deviation based on three to six independent biological replicates.

At the end of the ALE, we randomly selected six isolates from each of the three populations using the two carbon sources (36 colonies in total). We tested their ability to utilize C16OH and C20OH in growth experiments using the same carbon source from which they were obtained via ALE (Figure S2). From there, we selected the five best-performing adapted variants, C16OH-1 and C16OH-2 from C16OH, and C20OH-1, C20OH-2, and C20OH-3 from C20OH, and re-tested their phenotypes (Figure 2B–C). The growth rate and carrying capacity data show that the five adapted variants grow significantly faster than the wildtype (1.35x ±0.05 higher on C16OH, p<0.05, and 1.4x ±0.02 higher on C20OH, p<0.05) (Figure S3). Additionally, the specific growth rates of the adapted variants fell within the magnitude of the population from where they were isolated, indicating that the adapted variants well represent the population phenotype.

### 2.3 Mutation analysis sheds light on the molecular mechanisms behind optimized growth in fatty alcohols

To elucidate the genetic basis underlying the improved growth phenotypes, we performed whole-genome sequencing on the wild-type strain and the five adapted variants. To obtain a high-quality ancestral genome and facilitate the identification of plausible adaptive mutations in our isolated adapted variants, we re-sequenced the wild-type strain in our laboratory using a hybrid long- and short-read approach. The genome was assembled, annotated, and compared to the reference genome deposited on the NCBI repository (*GenBank*: AE015451.2) (Belda, 2016). Our wild-type genome and the reference genome differ only by 22 single-nucleotide polymorphisms (SNPs) and 27 short insertions or deletions (indels), as shown in Table S1, consistent with strain variation (Freddolino, 2012). The genomes of the adapted variants were obtained using deep short-read sequencing and compared with our wild-type genome. Mutation calling was performed using *breseq* (Deatherage & Barrick, 2014) and *snippy* (Seemann, 2015), both of which show the same mutation list, validating our results.

Two adapted variants, C20OH-2 and C20OH-3, contained only six and seven mutations, respectively. In contrast, C16OH-1, C16OH-2, and C20OH-1 each carried 30 to 38 mutations. These variants shared the same non-synonymous SNP, V466E (GTG to GAG) in the *mutL* gene, which encodes the DNA mismatch repair endonuclease, MutL (Modrich, 1991). In bacteria, MutL residues around 467-479 fall in the C-terminal endonuclease/metal-binding region that harbors the conserved DQHA(X)_2_E(X)_2_E motif required for MutL’s nicking activity. Alterations in this motif often attenuate mismatch repair and elevate mutation rates (i.e., hypermutation phenotype) as observed here (Correa, 2011).

Including those of the hypermutants, we detected a total of 91 mutations across the five adapted variants, including 63 SNPs and 28 short indels. Of these, 77 were located in coding regions and 14 in the intergenic areas. Of the 77 mutations in coding regions, 71 are non-synonymous mutations. These were located in 47 unique genes, indicating some levels of mutational convergence (Table S2).

To dissect the plausible contribution of mutations to the phenotype, we analyzed convergent mutations at the gene level across independent replicas, excluding synonymous mutations. Such analysis often highlights mutations that confer advantageous functions under the selection conditions of the experiment (Zhu, 2018; Mohamed, 2020; Ackermann, 2021). We found six mutations converging in the two C16OH-adapted variants, and two converging across the three C20OH-adapted variants (Table S3). Regardless of the carbon source from which they evolved, we identified eight convergent mutations (identical base) in at least three of the five isolates (Figure 3A, Table S3). Beyond convergence, we also analyzed the function of each mutated gene individually. The next sections describe the hypotheses and demonstrations we developed from these analyses.

**Figure 3.**
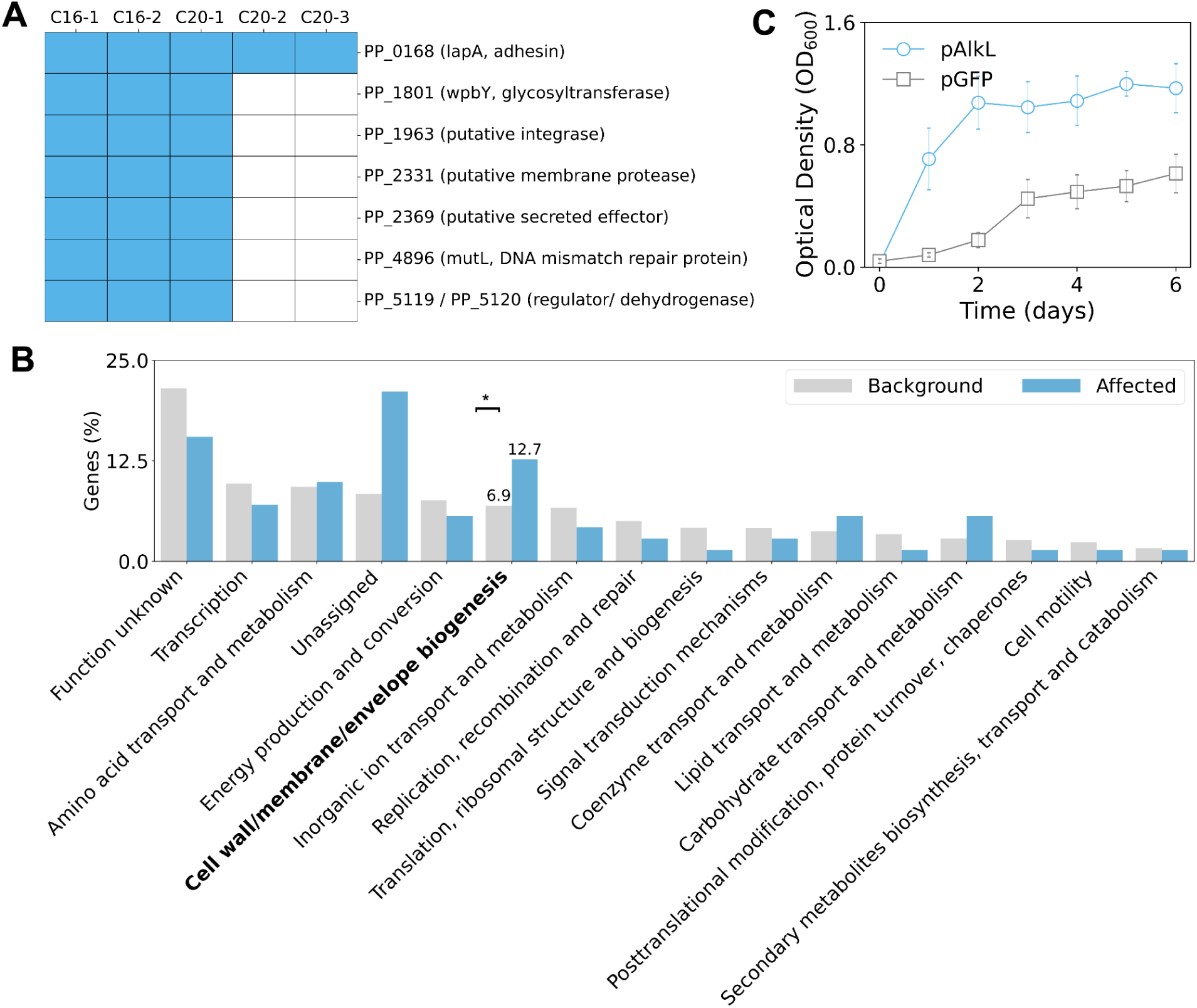
Mutational analysis points to a membrane-related adaptation mechanism. (A) Heatmap showing the occurrence of convergent non-synonymous mutations in C16OH and C20OH adapted variants. (B) Bar plot derived from the enrichment analysis by functional categories. COG categories which yielded zero occurrences in the affected genes, were omitted for visualization purposes. (C) Growth of wild-type *Pseudomonas putida* KT2440 expressing the heterologous AlkL transporter (pAlkL) or GFP as control (pGFP) on C16OH.

#### 2.3.1 Mutations in alcohol dehydrogenases are not essential to growth improvement

From the convergence analysis, we observed that the adapted variants C16OH-1, C16OH-2, and C20OH-1 harbor an SNP (G to A) in the intergenic region between genes PP_5119 (transcriptional regulator) and PP_5120 (aldehyde dehydrogenase). PP_5119 and PP_5120 are upstream of the genes PP_5121 and PP_5122, forming what appears to be a small operon (Figure S4A). Thus, we hypothesized that a mutation upstream of the transcriptional regulator (PP_5119) could affect the operon’s overall transcriptional level, revealing a functional role for these genes in fatty alcohol metabolism in *P. putida* KT2440. This is of particular interest because the genes PP_5121 and PP_5122 are homologous to the gene products of *laoB* (55% sequence identity) and *laoA* (70% sequence identity), respectively. LaoA and LaoB are the two subunits of an alcohol dehydrogenase, essential for growth in fatty alcohols ranging from 12 to 24 carbons in *Pseudomonas aeruginosa* PAO1 (Panasia, 2018).

We tested whether PP_5121 and PP_5122, homologs of *laoA* and *laoB,* were essential to growth in fatty alcohols in *P. putida* KT2440. We deleted the coding regions of PP_5121 and PP_5122 from the C16OH-1 and C20OH-1 adapted variants. The resulting strains, C16OH-1ΔPP_5121&2 and C20OH-1ΔPP_5121&2, grow in C16OH and C20OH, respectively, as well as the parental strains harboring these genes (Figure S4B–C), indicating that the LaoA and LaoB homologs are not essential for fatty alcohol utilization in *P. putida* KT2440.

As alcohol dehydrogenases are expected to be the first step and essential for alcohol metabolism, we looked for mutations in genes annotated for this function, regardless of convergence. We identified another SNP (G to A) in the adapted variant C16OH-1, located 93 base pairs upstream of the gene PP_4760, a predicted alcohol dehydrogenase. We again hypothesized that this gene would be essential for the phenotype. Similarly, the deletion of the coding region of PP_4760 from isolates C16OH-1 and C20OH-1 had no effect on the adapted variants’ capacity to utilize C16OH or C20OH (Figure S4B–C). Importantly, no other mutations were found in previously reported alcohol dehydrogenase genes, such as *adhP* (PP_3839), *pedE (*PP_2674*), and pedH* (PP_2679), which are known to mediate the oxidation of short-and medium-chain alcohols (Thompson, 2020). These results show that the alcohol dehydrogenase homologs, PP_5121 and PP_5122, and PP_4760 are not the only enzymes responsible for alcohol oxidation. As the *P. putida* KT2440 genome encodes 14 putative alcohol dehydrogenases, our data suggest alcohol oxidation is metabolically robust and not a limiting step under the tested conditions.

#### 2.3.2 Substrate uptake is the main bottleneck in fatty alcohol metabolism

We extended our analysis beyond catabolic genes to include all remaining mutations. In the convergence analysis, we observed four non-synonymous convergent changes occurring in membrane–associated genes across the adapted variants. These included: 1) *lapA* (PP_0168), mutated in all five variants, which encodes a surface adhesin, 2) *wpbY* (PP_1801), detected in three variants, encoding for a structural homolog of WaaG, a glycosyltransferase implicated in lipopolysaccharide biosynthesis in *P. aeruginosa, 3)* PP_2331, present in three variants, predicted to encode a membrane protease, and 4) PP_2369, also identified in three variants, annotated as a putative type III secreted effector-like protein (Scaletti, 2023; Espinosa-Urgel & Ramos-González 2023). Although these loci have distinct primary functions, together they point to modeling the outer surface and, thus, to cell–substrate interactions.

*P. putida* KT2440’s genome encodes 545 transporter genes (approximately 10% of its proteome). It is known that transporters can be promiscuous or multifunctional (Lewinson, 2006). Thus, it is tempting to speculate that different transporters could be mutated in different strains to achieve the same phenotype optimization. This genomic robustness may obscure clear patterns of gene-level convergence even when the phenotypic outcome is similar. This was indeed the case when we manually analyzed mutations regardless of convergence. For example, the isolate C16OH-1 exhibits non-synonymous mutations in PP_2656, a homolog of *ptsS* involved in cell adhesion in *P. aeruginosa* PAO1, and in PP_0880, which encodes an ABC transporter permease (Neznansky, 2014). C16OH-2 also harbors mutations in membrane-associated proteins, but different ones. We identified a non-synonymous SNP in PP_2754, which encodes a putative outer membrane porin of the OprD family, crucial for nutrient uptake in *P. aeruginosa PAO1* (Tamber, 2006).

To test whether adaptive mutations were preferentially found in envelope-associated proteins, we performed a functional category enrichment analysis on the mutational targets. We compiled a set by including all unique genes with non-synonymous coding mutations in all five adapted variants (47 genes). Also, if the mutation was found in an intergenic region, we included the two flanking genes to capture hypothetical regulatory effects on adjacent loci (26 genes). The total of 73 unique affected genes was assigned a single Clusters of Orthologous Groups (COG) category using eggNOG-mapper (Cantalapiedra, 2021). We compared the obtained gene set to the occurrence of all COG-assigned genes in wild-type *P. putida* KT2440 using two-sided Fisher’s exact tests with Benjamini-Hochberg correction (see *Methods*). This analysis revealed a significant enrichment of *cell wall/membrane/envelope biogenesis* COG category, consistent with envelope remodeling as a principal adaptive axis during growth on fatty alcohols (affected genes 12.7% vs. background genes 6.9%; Fisher’s exact q<0.05; Figure 3B).

To experimentally validate that membrane dynamics are important for fatty alcohol metabolism, we modified the wild-type strain to express the *alkL* gene from *Pseudomonas putida* GPo1. AlkL is a characterized transporter that facilitates the import of C7-C16 n-alkanes across the outer membrane (Grant, 2014; Grant, 2011). Previous studies have shown that it also facilitates the transport of medium-chain alcohols (C8OH to C12OH) (Lu, 2023). Growth assays were performed by comparing wild-type strains harboring either the *alkL*- or *gfp*-expressing plasmid. Remarkably, the expression of *alkL* significantly enhanced growth on C16OH (Figure 3C). The improvements matched the phenotypic gains observed in the C16OH-adapted variants, providing independent confirmation of the critical role of substrate uptake (Figure S5). As AlkL cannot accommodate substrates larger than C16, its overexpression did not enhance the growth of the wild-type strain in C20OH (Grant, 2014).

## 3. Discussion

In this study, we demonstrate that wild-type *Pseudomonas putida* KT2440 can utilize long-chain aliphatic alcohols (fatty alcohols) such as 1-hexadecanol (C16OH) and 1-eicosanol (C20OH) as the sole carbon source. While previous reports described *P. putida* KT2440 growth only on short- and medium-chain alcohols (up to C12OH) (Thompson, 2020; Lu, 2023), our results extend this substrate range to longer and less soluble alcohols relevant to the PE circular economy. Growth correlated strongly with substrate concentration, with high concentrations supporting faster growth rates and higher carrying capacities. This trend contrasts with that observed for medium-chain alcohol, such as 1-octanol (C8OH), where increased concentrations lead to toxicity rather than improved growth (Lu, 2023). This suggests that at longer chain lengths, toxicity is attenuated, likely due to reduced solubility, allowing *P. putida* KT2440 to exploit fatty alcohols when sufficient substrate is available in the media.

Five adaptive laboratory evolution passages on C16OH and C20OH revealed a rapid adaptive response, with improvements in growth kinetics after the first passage and stable performance thereafter. These results are in line with previous ALE studies in *P. putida* KT2440, where early adaptive sweeps also quickly shaped evolving populations, making population-level and clonal phenotypes very similar (Lim, 2021; Declerck, 2026). The fact that our adapted variants grew as fast as their parent populations suggests that beneficial mutations rapidly spread and reached fixation.

Through mutational analysis of the adapted variants compared with wildtype, we identified an intergenic mutation upstream of a small operon containing homologs of LaoAB, which are known to be essential for long-chain alcohol metabolism in *P. aeruginosa* PAO1 (Panasia, 2018), and a putative alcohol dehydrogenase (PP_4760). Although alcohol dehydrogenases should be essential for growth on fatty alcohols, deleting these genes in the adapted variants showed no observable impact on growth (Panasia, 2018). *P. putida* KT2440 encodes at least 14 alcohol dehydrogenases compared to 10 in *P. aeruginosa* PAO1 (Thompson, 2020). Such redundancy likely provides robustness (Fox, 1992; Wehrmann, 2017). Indeed, our data suggest that alcohol-oxidation capacity is not a limiting factor in fatty alcohol metabolism.

Moreover, mutations occurred consistently in genes involved in cell envelope and membrane functions. We identified four recurrent non-synonymous changes in membrane-associated genes. We also observed a significant enrichment of the *cell wall/membrane/envelope* biogenesis COG category among mutated genes, implicating envelope dynamics as a key adaptive axis during growth on fatty alcohols. Such mutations likely facilitated improved substrate access by altering surface hydrophobicity, permeability, or adhesion to hydrophobic substrates (Blesken, 2020). To experimentally validate the role of membrane dynamics and uptake in fatty alcohol utilization, we expressed AlkL from *P. putida* Gpo1 in the wild-type strain. AlkL is a characterized outer membrane transporter facilitating the import of medium-chain alkanes and alcohols (Grant, 2014; Grant, 2011). Its expression significantly enhanced growth on C16OH but not on C20OH, consistent with the known alkane-size limit of AlkL (up to C16). These results reinforce that substrate uptake capacity determines the upper limit of assimilable chain length, and that improved membrane permeability or transporter activity can directly enhance growth on fatty alcohols.

*P. putida* KT2440 encodes an unusually large repertoire of transporter genes (545 predicted, or ∼10% of its proteome) (Belda, 2016). To test whether any of these could act as an AlkL-like alkane transporter, we used Foldseek (Van Kempen, M. 2024) to compare the structure of AlkL against the predicted structures of all *P. putida* KT2440 proteins annotated as transporters. Yet, despite this extensive repertoire, neither the amino acid sequences nor the structures of *P. putida* KT2440’s transporters show significant identity to known systems for alkane assimilation. This suggests that either uncharacterized transporters mediate fatty alcohol uptake or that membrane remodeling collectively compensates for limited dedicated transport mechanisms.

## 4. Conclusions

Our study demonstrates that *Pseudomonas putida* KT2440 can utilize long-chain alcohols, such as 1-hexadecanol and 1-eicosanol, as a carbon source. It can rapidly evolve to efficiently assimilate these molecules, overcoming barriers related to their poor solubility. Through adaptive laboratory evolution, we generated isolates with improved growth phenotypes, supported by stable, heritable genetic changes. Genomic and functional analyses revealed that mutations in membrane-associated genes, including surface adhesion proteins and potential transporters drove these adaptations. These findings suggest that enhancing cell-substrate interactions is a key adaptive strategy for overcoming physical limitations to substrate uptake. Engineering the parental strain to express the *alkL* transporter from *P. putida* GPo1 reproduced the growth improvements on C16OH, providing experimental validation of the membrane transport hypothesis. Together, our results highlight the evolutionary plasticity of *P. putida* KT2440 and identify membrane remodeling as a promising target for future metabolic engineering strategies to valorize recalcitrant hydrophobic substrates.

## 5. Experimental procedures

### 5.1 Chemicals and culture media

All chemicals, including 1-hexadecanol and 1-eicosanol, as well as media components such as salts and trace elements, were purchased from Sigma-Aldrich (St. Louis, MO, USA). All reagents were of analytical grade and used without further purification. LB was obtained commercially (Difco™). AB minimal media was prepared with (NH_4_)_2_SO_4_ (2.0 g/L), Na_2_HPO_4_ (6.0 g/L), KH_2_PO_4_ (3.0 g/L), NaCl (3.0 g/L), and trace metals, CaCl_2_ (0.1 mM), MgCl_2_ (1.0 mM), and FeCl_3_ (3.0 *μ*M). All experiments were performed with sterile media in an autoclave.

### 5.2 Strains and plasmids

All strains and plasmids used in this study are listed in tables S4 and S5, respectively.

### 5.3 Growth experiments

*Pseudomonas putida* KT2440 was grown overnight on LB agar plates at 30 °C. Three individual colonies were picked and used to inoculate 4 mL of liquid LB medium, which was incubated overnight at 30 °C with shaking at 180 rpm. The resulting cultures were washed three times with AB minimal medium by centrifugation at 4,000 rpm for 4 minutes. Cell pellets were resuspended in AB minimal medium without a carbon source and adjusted to an optical density (OD₆₀₀) of 1.0. These cell suspensions were then used to inoculate 4 mL of AB medium supplemented with the corresponding carbon source at OD₆₀₀ of 0.05. Growth was monitored by OD₆₀₀ using a WPA CO 8000 Cell Density Meter (Biochrom Ltd. UK).

### 5.4 Adaptive laboratory evolution

Wild-type *Pseudomonas putida* KT2440 was grown overnight on LB agar plates at 30 °C. Three colonies were selected and grown overnight on 4-mL of LB media at 30 °C and 200 rpm. These cultures were then washed with AB minimal media and inoculated into 12-mL sterile glass tubes containing 4.0 mL of minimal media with 2.5 mM of 1-hexadecanol or 1-eicosanol as the sole carbon source and incubated at an initial OD_600_ of 0.05, 30 °C, 180 rpm. Adaptive Laboratory Evolution (ALE) was performed by transferring an aliquot of the previous culture into a sterile tube containing freshly prepared, sterile AB minimal media with the same carbon source concentration to an initial OD_600_ of 0.05. All transfers were made when the population reached the maximum optical density for at least 2 consecutive days. This process was repeated until the growth rate was stabilized for three consecutive passages. A glycerol (30% v/v) stock was created for each population at the end of the ALE experiment, which was stored at -80 °C.

### 5.5 Isolation of adapted variants

An LB agar plate was streaked from every final ALE population and incubated overnight at 30 °C. For each plate, three isolated colonies were randomly selected and grown overnight in 4-mL LB media, resulting in a total of 36 isolates. The next day a glycerol stock was created and stored at -80 °C. Additionally, these same glycerol stocks were used to streak fresh LB agar plates. These last plates belong to a single isolate that was used for the adapted variant selection. Three biological replicates from each isolate were tested in minimal media with the corresponding carbon source. Growth was followed by measuring optical density (OD_600_) every 24 hours. A single isolate from each population was selected for further analysis, giving rise to two adapted variants for C16OH and three for C20OH. The selection was based on growth rate and carrying capacity calculation based on custom python scripts. Finally, the optimized variants were subjected to whole-genome sequence analysis.

### 5.6 Molecular Biology

#### 5.6.1 Plasmid construction

All plasmids were constructed using the NEBuilder HiFi DNA Assembly system (New England Biolabs), a Gibson-type method that joins overlapping DNA fragments in a single reaction. DNA fragments were amplified by standard PCR techniques using the Q5 High-Fidelity DNA Polymerase (New England Biolabs), according to the manufacturer’s protocol. The oligonucleotide primers were designed accordingly to be used with the NEBuilder HiFi DNA Assembly system and synthesized by Sigma-Aldrich (Israel) (Table S6).

Non-replicative deletion plasmids (pDONRPEX18Gm) were generated by amplifying 500 bp regions upstream and downstream of each target gene from Pseudomonas putida KT2440 genomic DNA and cloning these flanking fragments into pDONRPEX18Gm.

The *alkL* gene encoding the medium-chain alkane transporter from *Pseudomonas putida* Gpo1 was chemically synthesized and cloned into pET28a(+) by Twist Biosciences. Such vector was used as template DNA for amplifying *alkL* and cloned into pS2313M, a broad-host-range plasmid carrying a kanamycin resistance cassette (KmR) and the pBBR1 origin of replication, in which the monomeric superfolder green fluorescent protein (*msfgfp*) reporter gene is placed under control of the constitutive synthetic promoter PEM7 by replacing the *msfgfp* gene sequence.

Assembly reactions were transformed into *E. coli* DH5ɑ by electroporation (25. kV, 4-5 ns) using a MicroPulser device (Bio-Rad Laboratories). Cell suspension was recovered with 900 uL of LB and incubated for two hours at 30 °C and 200 rpm and selected after plating in LB agar with the appropriate antibiotic marker (kanamycin 50 μg/mL and Gentamicin 10 μg/mL). All plasmid constructs were verified by sequencing performed by Plasmidsaurus, inc.

#### 5.6.2 Gene deletions

Gene deletions were performed using a two-step allelic exchange method based on homologous recombination (Hmelo, L. R, et al., 2015). The resulting pDONRPEX18Gm construct was introduced into *P. putida* KT2440 derivatives via conjugation with *E. coli* S17-1. Selection for the first homologous recombination event was performed on gentamicin media, resulting in chromosomal integration of the plasmid. Counterselection on sucrose-containing plates enabled the identification of double crossover events, leading to deletion of the target gene. Candidate mutants were screened and confirmed by PCR.

#### 5.6.3 Genetic engineering

A *P. putida* KT2440 WT single colony was picked from a previously inoculated LB agar plate and grown overnight in a four milliliter LB culture at 30 °C and 200 rpm. The cell pellet was recovered by centrifugation for four minutes and 4,000 rpm at room temperature. Then, it was washed twice with ice-cold, sterile 300 mM sucrose. The cell pellet was resuspended on 100 uL of the same solution. Such volume was mixed with 100 ng of either pS2313_AlkL or pS2313M and transferred to a sterile 1 mm electroporation cuvette. Plasmid DNA was incorporated by electroporation (25. kV, 4-5 ns) using a MicroPulser device (Bio-Rad Laboratories). Cell suspension was recovered with 900 uL of LB and incubated for two hours at 30 °C and 200 rpm. Selection of transformant colonies was made by plating and incubating overnight on LB agar plates supplemented with kanamycin (50 *μ*g/mL) at 30 °C.

### 5.7 DNA extraction and whole genome sequence analysis

The selected optimized variants were grown overnight in a 4-mL LB culture grown at 30 °C and 200 rpm. DNA was extracted using the *GenElute™ Bacterial Genomic DNA Kit* (Sigma-Aldrich). DNA sequencing was performed by SeqCenter (Pittsburgh, PA, USA). For all evolved variants, short-read Illumina sequencing was performed. Raw sequencing reads were quality-checked with FastQC and trimmed using Cutadapt (Martin, 2011). The resulting reads were aligned against the lab reference *P. putida* KT2440 WT to detect genetic variants *via breseq* and *snippy*. The affected genes were handled as indicated in the following section.

### 5.8 Enrichment analysis for functional categories

The reference genome was annotated using EggNOG-mapper, to assign functional categories (Cantalapiedra, 2021). We then evaluated whether affected genes were enriched or depleted in specific clusters of orthologous genes (COG) categories relative to the genome-wide background. For each COG category, we built a 2×2 contingency table that contrasts affected versus non-affected genes (i.e., the remainder of the annotated genome after removing affected genes), ensuring the two groups are disjoint. The odds ratio was computed with a Haldane–Anscombe correction of +0.5 to all cells when zeros occurred, and two-sided p-values were obtained from the hypergeometric tail probabilities (Fisher’s exact test). We excluded the *Unassigned* category from inferential testing but reported its counts for completeness; including it in the test is possible and does not affect the construction of the table. P-values across categories were adjusted for multiple comparisons using the Benjamini–Hochberg procedure.

## Acknowledgements

We thank the Standard European Vector Architecture (SEVA) collection for providing access to the plasmid, pS2313M, used in this study. We are also grateful to Prof. Nadav Kashtan (The Hebrew University of Jerusalem, Israel) for donating the wild-type *Pseudomonas putida* KT2440 strain.

## Author contributions

**RM**: Conceptualization, Data curation, Formal analysis, Investigation, Methodology, Software, Validation, Visualization, Writing - original draft and Writing, review and editing. **LN-G**: Conceptualization, Formal analysis, Funding acquisition, Investigation, Methodology, Project administration, Resources, Supervision, Validation, Visualization, Writing - original draft and Writing, review and editing.

