## Supplementary Information for "Adaptive laboratory evolution unlocks membrane permeability as a key limitation in long-chain alcohol metabolism by *Pseudomonas putida* KT2440"

Figure S1. Populations exhibit higher growth rates after five passages of adaptive laboratory evolution (ALE). 2

Figure S2. Isolated adapted variants show improved growth compared to wild-type (WT). 3

Figure S3. Growth rate and carrying capacity are higher in adapted strains than in wild-type strains. 4

Figure S4. Deletion of genes coding putative alcohol dehydrogenases does not impair growth on C16OH and C20OH. 5

Figure S5. Expression of the heterologous transporter AlkL enhances growth on C16OH. 6

Table S1. Mutations in the wild-type strain compared to the NCBI reference genome. Arrows indicate the direction of transcription of each gene relative to the genome. 7

Table S3. Occurrence of synonymous and non-synonymous convergent mutations in C16OH and C20OH isolates. 9

Table S4. Strains used in this work. 10

Table S5. Plasmids used in this work. 10

Table S6. Primers used in this work. 11

References 12

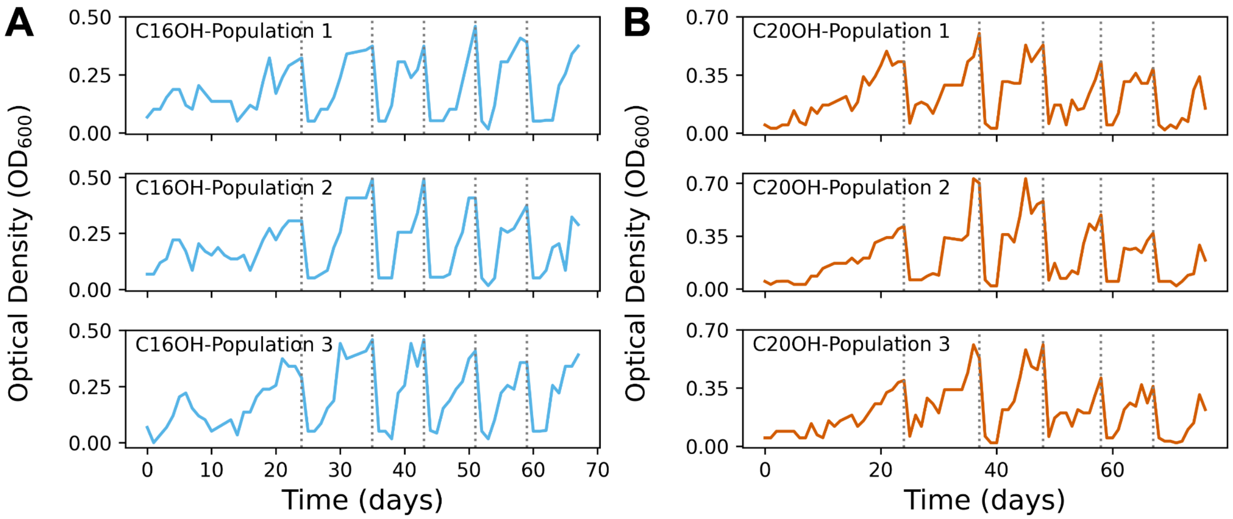


**Figure S1. Populations exhibit higher growth rates after five passages of adaptive laboratory evolution (ALE).**

Growth plots show the number of days required to reach maximum optical density for three independent populations of *Pseudomonas putida* KT2440 evolved on C16OH (A) and C20OH (B) across five serial passages. The passages are indicated by dashed lines.

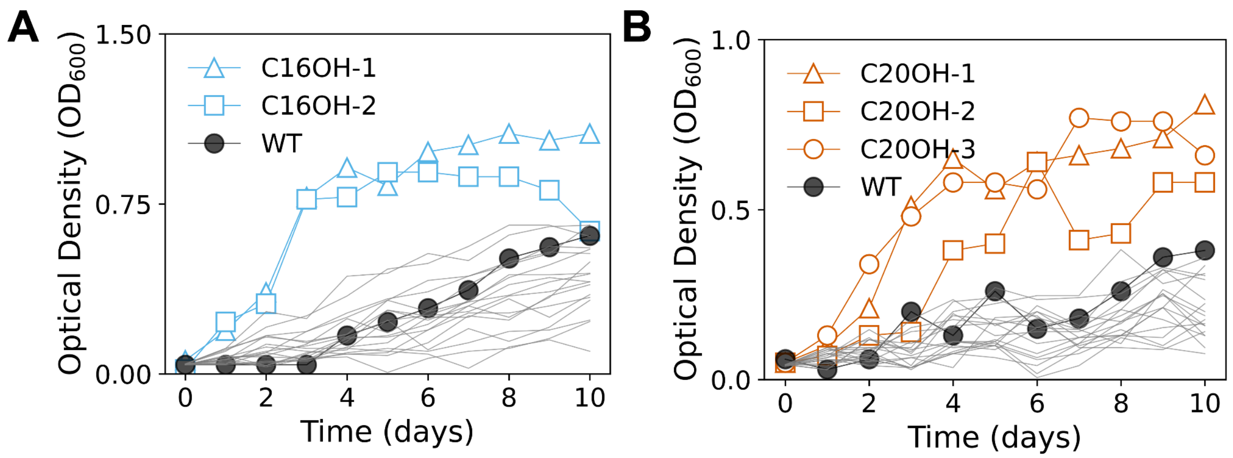


**Figure S2. Isolated adapted variants show improved growth compared to *wild-type* (WT).**

Selected adapted variants in C16OH (A) and C20OH (B) show markedly improved growth relative to wild-type (WT, black solid circles), while screened non-selected isolates are shown in gray.

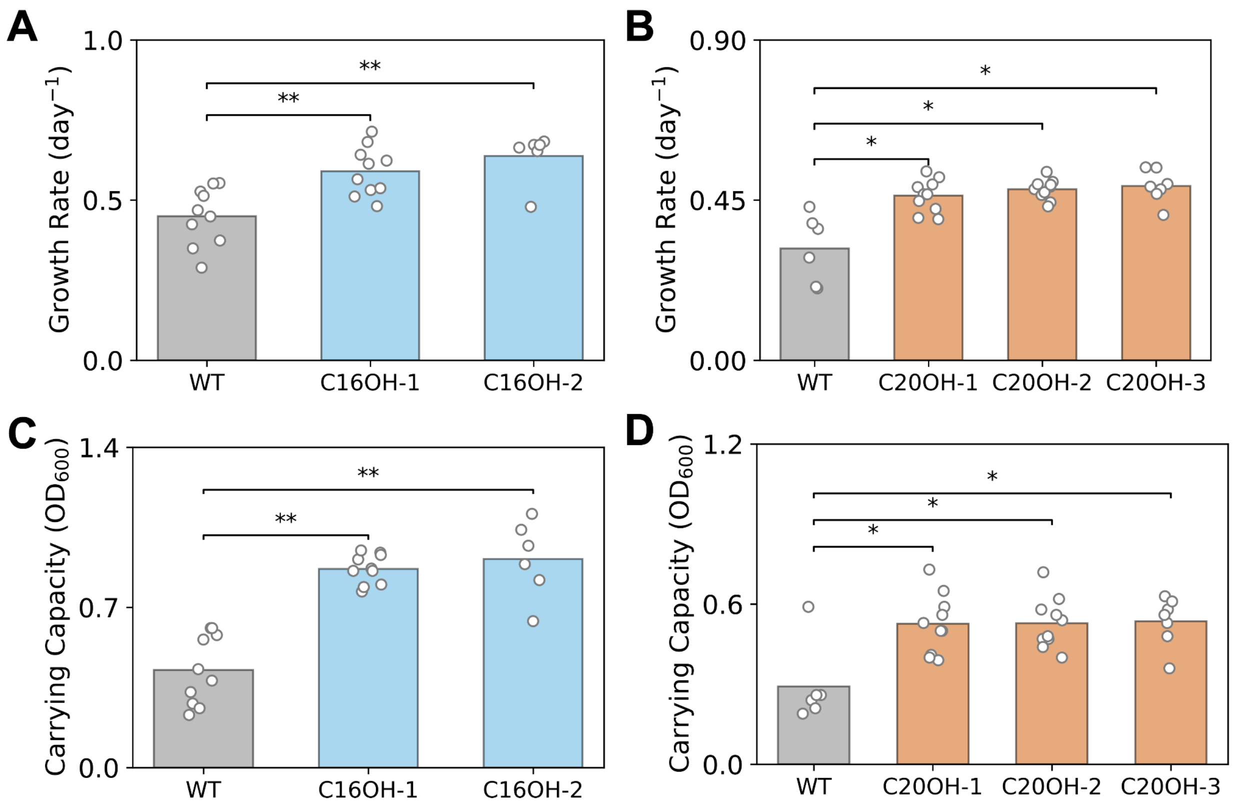


**Figure S3. Growth rate and carrying capacity are higher in adapted strains than in wild-type *strains*.**

Growth rate (day$^{-1}$) in C16OH (A) and C20OH (B), and carrying capacity ($OD_{600}$) in C16OH (C) and C20OH (D) of adapted variants and wild-type (WT). Bars represent mean values; dots represent individual six to ten biological replicates. Statistical comparisons were performed using unpaired two-sided t-tests. Stars indicate significance levels: $p < 0.05$ (*), $p < 0.01$ (**).

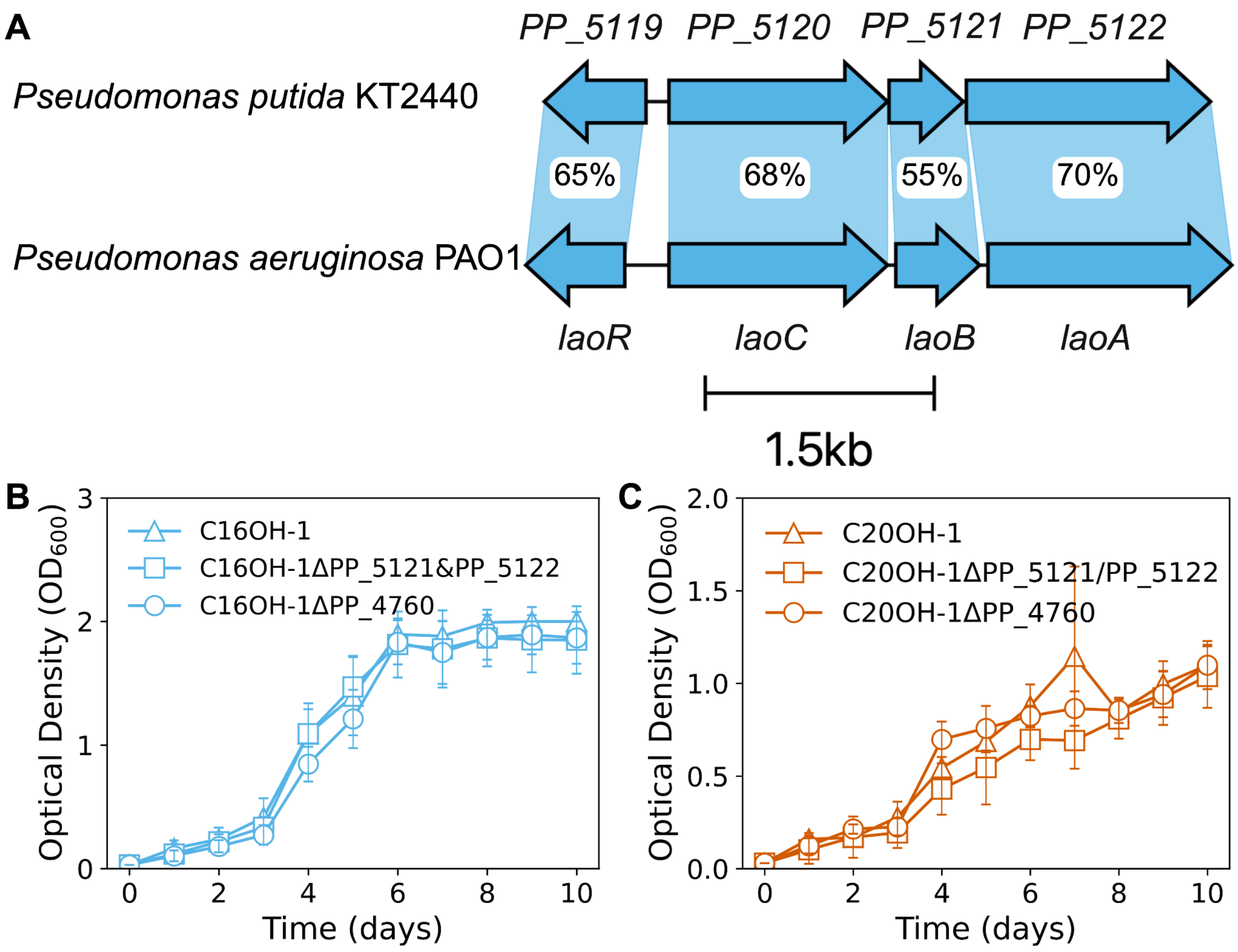


**Figure S4. Deletion of genes coding putative alcohol dehydrogenases does not impair growth on C16OH and C20OH.**

(A) Genomic organization of the homologous laoABCR locus in P. putida and P. aeruginosa PAO1. Arrows indicate the open reading frames and transcriptional orientation. Percentages denote amino-acid sequence identity. Growth assays of evolved isolates, C16OH-1 and C20OH-1, and its derivatives lacking candidate genes in C16OH (B) and C20OH (C) as sole carbon sources.

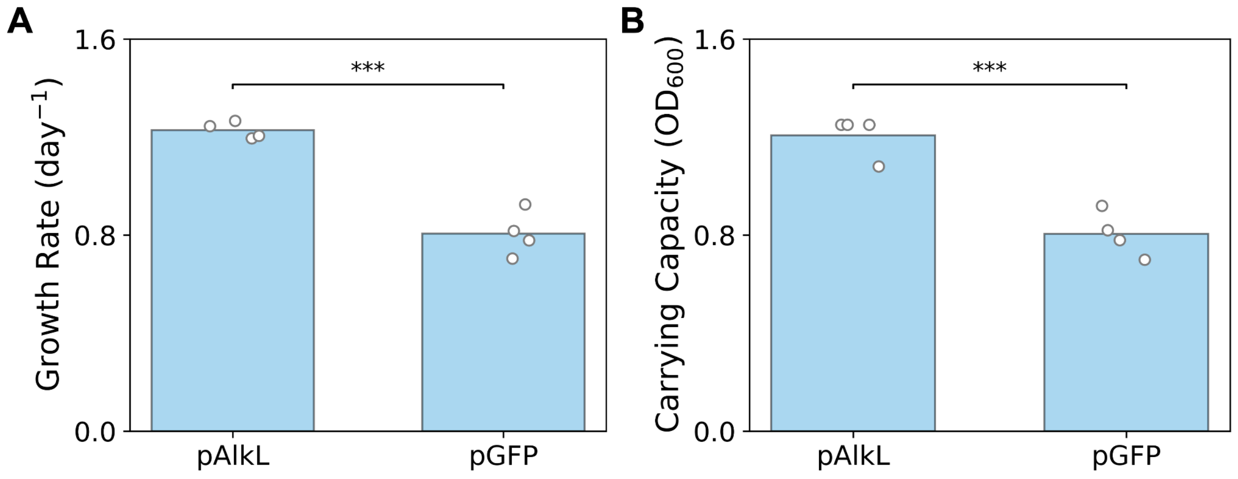


**Figure S5. Expression of the heterologous transporter AlkL enhances growth on C16OH.** (A) Growth rates of *wild-type P. putida* KT2440 carrying pAlkL or pGFP. (B) Corresponding carrying capacities. Bars show means of four individual biological replicates (shown as dots). Stars indicate significance levels: $p$ < 0.01 (***)

**Table S1. Mutations in the wild-type strain compared to the NCBI reference genome. Arrows indicate the direction of transcription of each gene relative to the genome.**

| Mutation | Locus Tag | Annotation |
|---|---|---|
| +A | *PP_0224 ← / ← sctC* | Monooxygenase/ ABC transporter |
| +C | *hslO → / → PP_0254* | chaperonin/unknown function |
| +T | *PP_0278 →* | unknown function |
| A→G | *hisF → / ← cbcV* | phosphate synthase/ABC transporter |
| A→G | *uhpA → / → PP_0411* | regulator UhpA/ABC transporter |
| +C | *tdcG-II ← / ← gcvP-I* | L-serine dehydratase/glycine dehydrogenase |
| +C | *PP_1242 ← / ← PP_1244* | unknown function/unknown function |
| 2 bp→AC | *trpS → / → zapE* | tRNA ligase/ nucleoside triphosphate hydrolase |
| 1 bp→CG | *trpS → / → zapE* | tRNA ligase/ nucleoside triphosphate hydrolase |
| +C | *trpS → / → zapE* | tRNA ligase/ nucleoside triphosphate hydrolase |
| +C | *rluA ← / ← minE* | rRNA and tRNA synthase/Cell division factor |
| C→T | *PP_2589 →* | Aldehyde dehydrogenase family protein |
| +G | *PP_3394 ← / ← PP_3395* | CoA lyase/Transcriptional regulator |
| T→C | *PP_3486 ← / ← PP_3487* | cytochrome c/unknown function |
| A→T | *PP_3486 ← / ← PP_3487* | cytochrome c/unknown function |
| 2 bp→TC | *PP_4061 → / ← PP_4063* | unknown function/-CoA ligase |
| +G | *PP_4061 → / ← PP_4063* | unknown function/-CoA ligase |
| +C | *PP_4061 → / ← PP_4063* | unknown function/-CoA ligase |
| $(T)_{7\rightarrow6}$ | *gltA → / ← yeiW* | citrate synthase/putative oxidoreductase |
| T→G | *gltA → / ← yeiW* | citrate synthase/putative oxidoreductase |
| 2 bp→CT | *gltA → / ← yeiW* | citrate synthase/putative oxidoreductase |
| +C | *gltA → / ← yeiW* | citrate synthase/putative oxidoreductase |
| A→G | *gltA → / ← yeiW* | citrate synthase/putative oxidoreductase |
| C→T | *gltA → / ← yeiW* | citrate synthase/putative oxidoreductase |

| | | |
|---|---|---|
| C→G | *gltA* → / ← *yeiW* | citrate synthase/putative oxidoreductase |
| 2 bp→CT | *gltA* → / ← *yeiW* | citrate synthase/putative oxidoreductase |
| Δ1 bp | *gltA* → / ← *yeiW* | citrate synthase/putative oxidoreductase |
| +T | *gltA* → / ← *yeiW* | citrate synthase/putative oxidoreductase |
| A→G | *gltA* → / ← *yeiW* | citrate synthase/putative oxidoreductase |
| A→C | *gltA* → / ← *yeiW* | citrate synthase/putative oxidoreductase |
| Δ1 bp | *gltA* → / ← *yeiW* | citrate synthase/putative oxidoreductase |
| A→G | *gltA* → / ← *yeiW* | citrate synthase/putative oxidoreductase |
| A→C | *gltA* → / ← *yeiW* | citrate synthase/putative oxidoreductase |
| G→C | *gltA* → / ← *yeiW* | citrate synthase/putative oxidoreductase |
| C→T | *gltA* → / ← *yeiW* | citrate synthase/putative oxidoreductase |
| +T | *gltA* → / ← *yeiW* | citrate synthase/putative oxidoreductase |
| C→G | *gltA* → / ← *yeiW* | citrate synthase/putative oxidoreductase |
| C→T | *gltA* → / ← *yeiW* | citrate synthase/putative oxidoreductase |
| C→G | *gltA* → / ← *yeiW* | citrate synthase/putative oxidoreductase |
| A→C | *gltA* → / ← *yeiW* | citrate synthase/putative oxidoreductase |
| C→T | *gltA* → / ← *yeiW* | citrate synthase/putative oxidoreductase |
| +GGC | *PP_4387* ← / ← *flgE* | unknown function/Flagellar hook protein |
| +GCC | *PP_4887* ← / ← *PP_4888* | unknown function/chemotaxis transducer |
| +G | *rhlE-II* → / ← *yceI* | RNA helicase/ stress response factor |
| G→C | *rhlE-II* → / ← *yceI* | RNA helicase/ stress response factor |
| +CGGG | *PP_4986* ← / ← *PP_4987* | Channel protein/Chemotaxis protein |
| T→C | *xpt* → / ← *PP_5266* | phosphoribosyltransferase/ acetyl-CoA hydrolase |

**Table S3. Occurrence of synonymous and non-synonymous convergent mutations in C16OH and C20OH isolates.**

| Mutation | Locus Tag | Annotation | Occurrence in variants | | | | |
|---|---|---|---|---|---|---|---|
| | | | C16OH -1 | C16OH -2 | C20OH -1 | C20OH -2 | C20OH -3 |
| C→T | *PP_0168* | S4900S (AGC→AGT) | Yes | Yes | Yes | Yes | Yes |
| G→A | | Q4908Q (CAG→CAA) | Yes | Yes | Yes | Yes | Yes |
| T→C | | G4909G (GGT→GGC) | Yes | Yes | Yes | Yes | Yes |
| C→T | | V3353V (GTC→GTT) | Yes | Yes | Yes | Yes | Yes |
| C→A | | Q3355N (CAG→AAC) | - | Yes | Yes | Yes | Yes |
| G→C | | Q3355N (CAG→AAC) | - | Yes | Yes | Yes | Yes |
| $(C)_{5\rightarrow4}$ | *PP_1801* | coding (906/1143 nt) | Yes | Yes | Yes | - | - |
| G→T | *PP_1963* | A526S (GCC→TCC) | Yes | Yes | Yes | - | - |
| T→C | *PP_2331* | V468A (GTG→GCG) | Yes | Yes | Yes | - | - |
| A→G | *PP_2369* | F263L (TTC→CTC) | Yes | Yes | Yes | - | - |
| A→T | *PP_4896* | V466E (GTG→GAG) | Yes | Yes | Yes | - | - |
| G→A | *PP_5119 / PP_5120* | intergenic (-116/-35) | Yes | Yes | Yes | - | - |

**Table S4. Strains used in this work.**

| Name | Description | Reference |
|---|---|---|
| KT2440 | *wild-type Pseudomonas putida* KT2440 strain | Bagdasarian, 1981 |
| C16OH-1 | 1-hexadecanol ALE-derived *P. putida* KT2440 variant 1. | This work |
| C16OH-2 | 1-hexadecanol ALE-derived *P. putida* KT2440 variant 2. | This work |
| C20OH-1 | 1-eicosanol ALE-derived *P. putida* KT2440 variant 1. | This work |
| C20OH-2 | 1-eicosanol ALE-derived *P. putida* KT2440 variant 2. | This work |
| C20OH-3 | 1-eicosanol ALE-derived *P. putida* KT2440 variant 3. | This work |
| pGFP | *P. putida* KT2440 harboring plasmid pS2313M | This work |
| pAlkL | *P. putida* KT2440 harboring plasmid pS2313_AlkL | This work. |
| DH5$\alpha$ | *Escherichia coli* cloning host: $F^{-}$ $\lambda^{-}$ *endA1 glnX44(AS) thiE1 recA1 relA1 spoT1 gyrA96*($Nal^{R}$) *rfbC1 feoR nupG* $\Phi$*80(ΔlacZM15) Δ(argF-lac)U169 hsdR17* ($r_K^- m_K^+$) | Hanahan & Meselson, 1983 |
| S17-1 | *Escherichia coli* conjugation donor strain: thi pro hsdR *recA* RP4-2-Tc::Mu-Km::Tn7 integrated into chromosome | Simonl, 1983 |

**Table S5. Plasmids used in this work.**

| Name | Description | Reference |
|---|---|---|
| pS2313M | $Km^R$, *ori pBBR1*, $P_{EM7}$, *msfgfp* | Volke, 2020 |
| pS2313_AlkL | $Km^R$, *ori pBBR1*, $P_{EM7}$, *alkL* | This work. |
| pDONRPEX18Gm | Suicide vector: $Gm^R$, *sacB*$^+$, $ori_{TRP4}$, Gateway-compatible | Hmelo, 2015 |

**Table S6. Primers used in this work.**

| Name | Sequence 5' to 3' |
| --- | --- |
| pDONR_rev | *cgactctagaggatccccgggtac* |
| pDONR_for | *catcatgaaagcttggcactggcc* |
| ups_4760_for | tgcatgcctgcaggtcgacttgccggctgggtattctgacc |
| ups_4760_rev | cctcacttcacatgggtgggtctccgtatcagagc |
| ds_4760_for | cccacccatgtgaagtgagggcagggtagctg |
| ds_4760_rev | cggtacccggggatcctctaggcgaatgtgcgcgatctgct |
| up_laoAB_rev | aaatagcagacatacgaatgtcatgcgctgtgtgccaggcgg |
| laoAB_for | aagcttaggaggaaaaacatatgcaccgccgcgacctgctg |
| ups_laoAB_for | ccggggatcctctagagtcgcaaaggcgcgcaagtactcgactt |
| ups_laoAB_rev | gtgccaagctttcatgatgtcgaggaaatgaacgaataacgctccgacg |
| pS2313M_rev | atgtttttcctcctaagcttgcatgcctg |
| pS2313M_for | cattcgtatgtctgctatttcgcgtatagaactagtcttg |
| AlkL_for | aagcttaggaggaaaaacatATGAGTTTTTCTAATTATAAAGTAATCGCG |
| AlkL_rev | aaatagcagacatacgaatgTTACTAGAAAACATATGACGCACCAA |
